# Serological and metagenomic interrogation of cerebrospinal fluid implicates enteroviruses in pediatric acute flaccid myelitis

**DOI:** 10.1101/666230

**Authors:** Ryan D. Schubert, Isobel Hawes, Prashanth S. Ramachandran, Akshaya Ramesh, Emily D. Crawford, John E. Pak, Wesley Wu, Carly K. Cheung, Brian D. O’Donovan, Cristina M. Tato, Amy Lyden, Michelle Tan, Rene Sit, Gavin Sowa, Hannah A. Sample, Kelsey C. Zorn, Debarko Banerji, Lillian M. Khan, Riley Bove, Stephen L. Hauser, Amy A. Gelfand, Bethany Johnson-Kerner, Kendall Nash, Kalpathy S. Krishnamoorthy, Tanuja Chitnis, Joy Z. Ding, Hugh J. McMillan, Charles Y. Chiu, Benjamin Briggs, Carol A. Glaser, Cynthia Yen, Victoria Chu, Debra A. Wadford, Samuel R. Dominguez, Terry Fei Fan Ng, Rachel L. Marine, Adriana S. Lopez, W. Allan Nix, Ariane Soldatos, Mark P. Gorman, Leslie Benson, Kevin Messacar, Jennifer L. Konopka-Anstadt, M. Steven Oberste, Joseph L. DeRisi, Michael R. Wilson

## Abstract

**Background:** Since 2014, the United States has experienced a biennial spike in pediatric acute flaccid myelitis (AFM). Epidemiologic evidence suggests non-polio enteroviruses (EVs) are a potential etiology, yet EV RNA is rarely detected in cerebrospinal fluid (CSF) and only inconsistently identified from the respiratory tract, serum, or stool.

**Methods:** We interrogated CSF from children with AFM (n=42) and pediatric controls with other neurologic diseases (OND) (n=58). Samples were incubated with T7 bacteriophage expressing 481,966 sixty-two amino acid peptides with a fourteen amino acid overlap tiled across all known vertebrate virus and arbovirus genomes, an adaption of the VirScan method. Antibody-bound phage were deep sequenced to quantify enriched peptides with normalized counts expressed as reads per hundred thousand (rpK). EV antibody findings were confirmed with ELISA using whole viral protein 1 (VP1) from contemporary enterovirus (EV) A71 and D68 strains. Separately, metagenomic next-generation sequencing (mNGS) of CSF RNA, both unbiased and with targeted enrichment for EVs, was performed.

**Results:** The most significantly enriched viral family by VirScan of CSF in AFM versus OND controls was *Picornaviridae* (mean rpK 11,266 versus mean rpK 950, p-adjusted < 0.001, Wilcoxon signed-rank test with Bonferroni adjustment). Enriched *Picornaviridae* peptides belonged almost entirely to the genus *Enterovirus.* The mean EV VP1 ELISA signal in AFM (mean OD 0.51) was significantly higher than OND controls (mean OD 0.08, p-value < 0.001, Mann-Whitney test). mNGS did not detect additional enterovirus RNA in CSF.

**Conclusion:** Despite the rare detection of EV RNA in the CNS of patients with AFM, a pan-viral serologic assay identified high levels of CSF EV antibodies in AFM CSF compared to CSF from OND controls. These results provide further evidence for a causal role of non-polio enteroviruses in AFM.

## Introduction

First detected in California in 2012, the United States has experienced seasonal, biennial increases in the incidence of acute flaccid myelitis (AFM) cases.^1^ Since 2014, the Centers for Disease Control and Prevention (CDC) has reported over 500 confirmed cases.^2–6^ The nationwide surges in AFM in 2014, 2016, and 2018 have coincided temporally and geographically with outbreaks of enterovirus (EV) D68 and EV-A71 infections.^3,7–10^ EVs, including poliovirus, are well recognized for their neuroinvasive capacity and resultant central nervous system (CNS) pathology, ranging from self-resolving aseptic meningitis to fulminant, sometimes fatal, brainstem encephalitis, and to myelitis leading to permanent debilitating paralysis.^11^

Despite the temporal association between EV-D68 and EV-A71 outbreaks and AFM and a mouse model that recapitulates the AFM phenotype with a contemporary EV-D68 strain,^12^ the etiology of AFM has been difficult to confirm.^13,14^ Thus, concerns persist that AFM could result from yet-to-be-identified pathogens or a para-infectious immune response. This is due, in part, to the fact that less than half of children with AFM have had EV detected in a non-sterile biologic specimen (nasopharyngeal or oropharyngeal swabs most commonly, rectal and stool samples less commonly), and no other alternative candidate etiologic agents have been identified in the remaining children.^4^ In addition, only 2% of children with AFM have EV nucleic acid detected in cerebrospinal fluid (CSF).^15,16^

The immune privileged status of the CNS makes direct detection of viral nucleic acid or indirect discovery of intrathecal anti-viral antibodies an important step in linking a pathogen to a neuroinfectious disease. We interrogated CSF from AFM patients (n=42) from recent outbreaks with unbiased ultra-deep metagenomic next-generation sequencing (mNGS), including with a novel CRISPR-Cas9 based enrichment technique. Furthermore, to search for virome-wide antibody signals that might be associated with AFM, we employed the VirScan approach that was previously developed to detect antibodies to all known human viruses.^17^ To improve upon this detection method we generated a large and more finely tiled peptide library in the T7 phage display vector described in detail in Methods.

## Methods

Detailed methods for data collection, human subjects review, mNGS, VirScan, bioinformatics, and independent confirmatory testing with ELISA are provided in the Supplemental Appendix.

### Case-Control Design

All AFM cases met the 2018 US Council of State and Territorial Epidemiologists case definition for probable or confirmed AFM (Supplemental Table 1).^18^ Patient samples were collected either through enrollment in research studies or through public health surveillance. In addition, residual banked CSF was obtained from children with other neurologic diseases (ONDs) without suspected primary EV infection for controls.

### Metagenomic Sequencing Library Preparation

RNA sequencing libraries were prepared using a previously described protocol optimized and adapted for miniaturization and automation.^19^ Libraries were sequenced on a NovaSeq 6000 machine (Illumina) to generate 150 nucleotide (nt), paired-end reads. Samples were also sequenced after enrichment for EV-A71 and EV-D68 genomes using FLASH (Finding Low Abundance Sequences by Hybridization, Supplemental Table 2).^20^ All NGS libraries were depleted of host ribosomal RNA with DASH and spiked-in with External RNA Controls Consortium (ERCC) sequences as previously described.^21,22^

### Metagenomic Bioinformatics

As previously described, pathogens were identified from raw mNGS sequencing reads using IDseq v3.2, a cloud-based, open-source bioinformatics platform designed for detection of microbes from mNGS data.^23^

### Pan-Viral CSF Serologic Testing with VirScan

The previously published VirScan method is a variation on phage immunoprecipitation-sequencing (PhIP-Seq), displaying viral peptides on the outer surface of bacteriophage for the purposes of antibody detection followed by deep sequencing.^23,24^ Similarly, we constructed a T7 bacteriophage display library comprised of 481,966 sixty-two amino acid peptides with a 14 amino acid overlap tiled across a representative set of full-length, vertebrate, mosquito-borne, and tick-borne viral genomes downloaded from the UniProt and RefSeq databases in February 2017. After amplification, phage libraries were incubated with 2 μL of patient CSF overnight and then immunoprecipitated for two rounds. Barcoded phage DNA was sequenced on a HiSeq 4000 machine (Illumina) using 150 nt paired-end reads.

### VirScan Bioinformatics

Sequencing reads were aligned to a reference database comprising the full viral peptide library. Peptide counts were normalized by dividing by the sum of counts and multiplying by 100,000 (reads per hundred thousand (rpK)).^25–27^ Phage results were filtered using a cutoff fold-change of greater than 10 above the mean background rpK generated from null IPs.

### Independent Validation with ELISA

To independently validate our VirScan results, we generated recombinant viral protein 1 (VP1) from recent AFM-associated EV-A71 and EV-D68 strains and performed ELISA to detect EV antibodies with AFM CSF samples for which sufficient CSF remained (n=26) and OND controls (n=50). Signal was measured as the optical density (OD) at 450 nm. For each sample, we considered the higher of the two (EV-A71 or EV-D68) OD values when analyzing cases and controls.

## Results

### Cases and Controls

42 AFM cases and 58 OND controls were included in the study (Supplemental Figure 1). Patient demographics are described in Table 1. The AFM cases were younger (median age 37.8 months, interquartile range [IQR], 11 to 64 months) than the OND controls (median age 120 months, IQR, 66 to 174 months), with a p-value of 0.0497 (as determined by an unpaired parametric t-test). There was a higher proportion of males in the AFM cases. AFM cases and OND controls from the Western and Northeastern USA (Supplemental Figure 2) make up the majority of both categories. Cases from 2018 make up the majority of the AFM cases.

**Table 1.** Characteristics of the Patients at Baseline.

|  | AFM Cases | OND Controls |
| --- | --- | --- |
| N | 42 | 58 |
| Age – median (IQR), months | 38 (11 to 64) | 120 (66-174) |
| Sex – no. (%) |  |  |
| Female | 13 (31) | 32 (55) |
| Male | 29 (69) | 26 (45) |
| Region – no. (%) |  |  |
| United States |  |  |
| West | 20 (48) | 37 (64) |
| South | 7 (17) | 4 (7) |
| Midwest | 3 (7) | 4 (7) |
| Northeast | 11 (26) | 9 (16) |
| International |  |  |
| South America | 0 (0) | 2 (3) |
| Canada | 1 (2) | 0 (0) |
| North Atlantic Island | 0 (0) | 1 (2) |
| Middle East | 0 (0) | 1 (2) |
| Year – no. (%) |  |  |
| 2014 | 5 (12) | 10 (17) |
| 2015 | 0 (0) | 14 (24) |
| 2016 | 2 (5) | 12 (21) |
| 2017 | 0 (0) | 8 (14) |
| 2018 | 34 (81) | 14 (24) |
| Season – no. (%) |  |  |
| Spring | 1 (2) | 18 (31) |
| Summer | 12 (29) | 8 (14) |
| Fall | 24 (57) | 20 (34) |
| Winter | 5 (12) | 12 (21) |
| Suspected Etiology – no. (%) |  |  |
| Infectious | - | 23 (40) |
| Autoimmune | - | 22 (38) |
| Non-inflammatory | - | 6 (10) |
| Malignancy | - | 3 (5) |
| Unavailable | - | 4 (7) |

NB: Percentages may not total 100 because of rounding.

### Ultra-Deep Metagenomic Next-Generation Sequencing Rarely Detects Enterovirus in AFM

We obtained an average of 433 million 150 nt paired-end reads per sample (range, 304 - 569 million reads per sample). Based on the ERCC RNA spike-ins, we estimated that our mean limit of detection was 5.48 attograms (range, 3.92 to 17.47 attograms).^21^ EV-A71 was detected in one AFM sample at 71.31 rpM (1497.3 rpM in FLASH-NGS, Supplemental Table 3). This sample was previously known to be EV-A71 positive by Sanger sequencing. No other pathogenic organisms were detected in this or any of the other AFM samples.

### CSF VirScan Testing Detects Enterovirus in AFM

The most significantly enriched viral family by VirScan of CSF in AFM cases (n = 42) versus OND controls (n = 58) was *Picornaviridae* (mean rpK 11,266, IQR 16,324 versus mean rpK 950 IQR 948, p-adjusted = 1 x 10^−7^ Wilcoxon signed-rank test with Bonferroni adjustment, Supplemental Table 4). Enriched *Picornaviridae* peptides belonged almost entirely to the genus *Enterovirus* (Figure 1A-C, Supplemental Table 5). Peptides mapping to *Caliciviridae*, *Ascoviridae*, *Baculoviridae* and unclassified viruses were also significantly enriched in AFM relative to OND controls (p-adjusted = 0.004, 0.025, 0.021, and 0.039, respectively by Wilcoxon signed-rank test with Bonferroni adjustment) but with a mean rpK 6.4 to 74.7 times lower than for *Picornaviridae* (Figure 1A and Supplemental Table 4). We detected 3.4-fold more *Caliciviridae* in AFM versus OND (mean rpK 151 IQR 132 versus mean rpK 44 IQR 0, p-adjusted < 0.01). We did not further consider viruses with non-human hosts.

**Figure 1.**
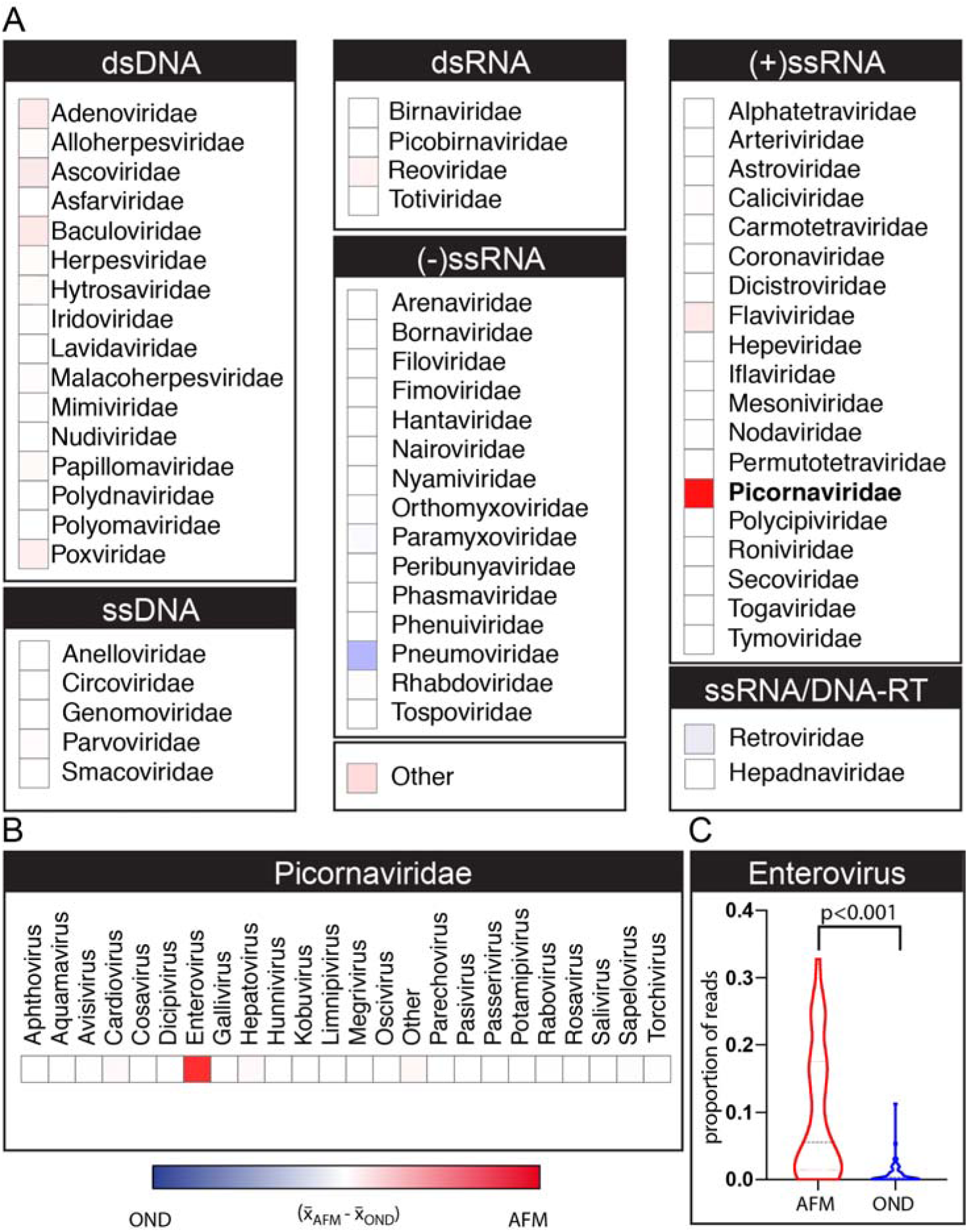
Enterovirus Immunoreactivity in Acute Flaccid Myelitis on a Pan-Viral Phage Display Assay. **(A)** Viral families detected by VirScan or phage display immunoprecipitation with next-generation sequencing (PhIP-Seq) sorted by their Baltimore classification. Heatmap color intensity was calculated by subtracting the mean reads per hundred thousand sequenced (rpK) in the other neurologic disease (OND) cerebrospinal fluid sample set (n=58) from that observed in acute flaccid myelitis (AFM) CSF (n=42). The maximum and minimum color intensities reflect +11,000 and −11,000 rpK, respectively. The strongest intensity is observed in the *Picornaviridae* family (boldface type). **(B)** Genus *Enterovirus* demonstrating the strongest enrichment in family *Picornaviridae*. **(C)** Violin plot of the proportion of *Enterovirus* phage per patient with mean and first and third quartile indicated by horizontal lines; Mann-Whitney test corrected for multiple comparisons with a Bonferroni adjustment.

Enriched EV peptides were derived from proteins across the EV genome (Figure 2A, Supplemental Table 6). Among capsid protein sequences, KVPALQAAEIGA in VP1 has been previously reported to be an immunodominant linear EV epitope.^28^ Peptides containing this and related overlapping epitopes were enriched in our data across AFM patients, with multiple sequence alignment revealing a consensus motif of PxLxAxExG (Figure 2B). Another immunodominant epitope was to a conserved, linear portion of 3D^pol^ (Figure 2C).

**Figure 2.**
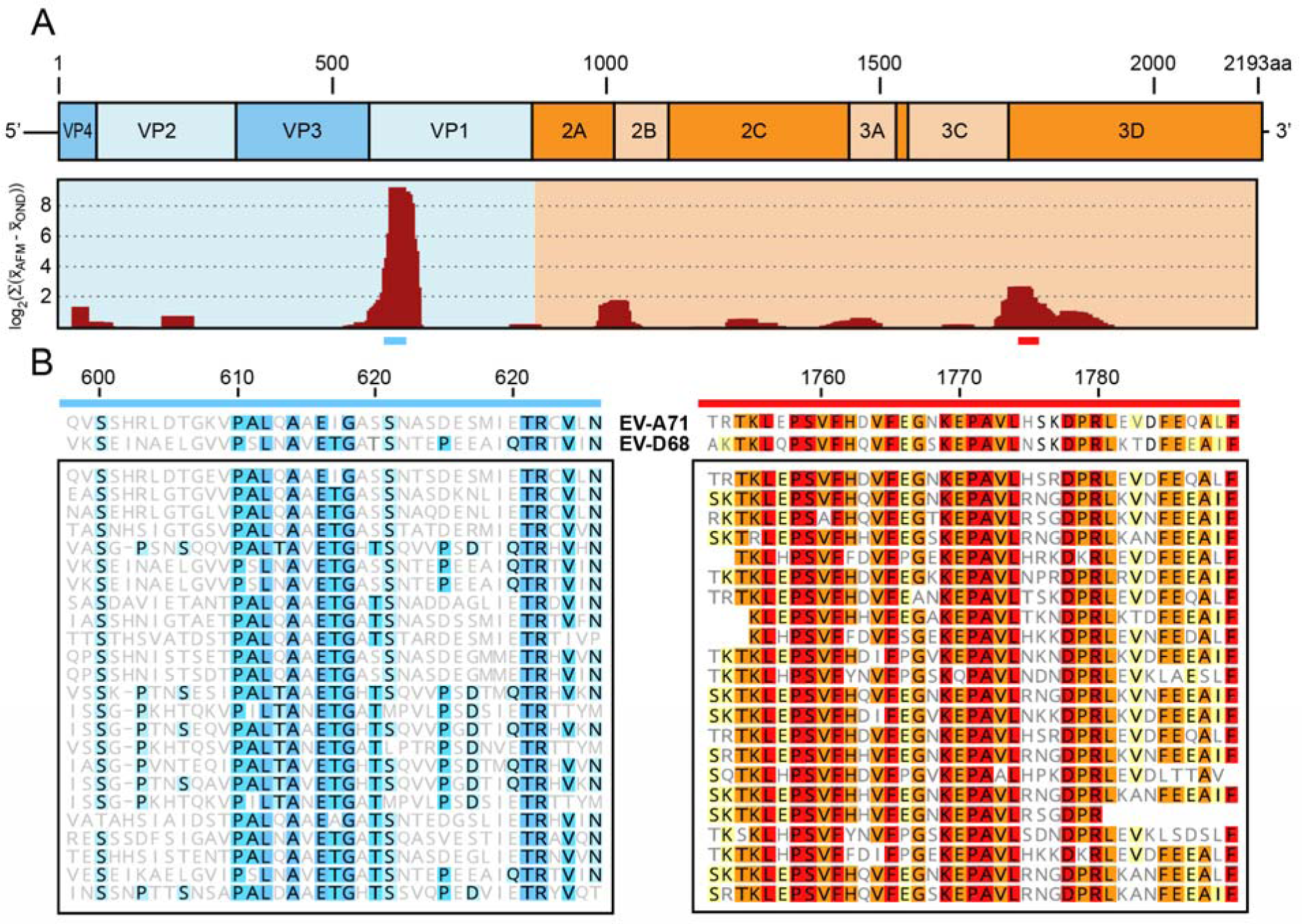
Primary Enterovirus Antigens Identified by Pan-Viral Phage Display in Acute Flaccid Myelitis. We identified 263 unique, enriched antigens with taxonomic identifications mapping to enterovirus (EV) across all acute flaccid myelitis (AFM) cerebrospinal fluid samples (n=42). **(A)** 252 of 263 EV derived peptides were mapped by BLASTP to the 2,193 amino acid polyprotein of EV-A71 (Genbank Accession AXK59213.1) as a model reference. The relative recovery of these peptides by VirScan is plotted as the log2 of the sum of the differences in the mean signal generated in the AFM and other neurologic disease (OND) cohorts, using a moving average of 32 amino acids, advanced by 4 amino acid steps. **(B)** Multiple sequence alignment of representative enriched EV-derived peptides for the VP1 (blue bar) and 3D (red bar) proteins. Sequences from EV-D68 (Genbank Accession AIT52326.1) and EV-A71 (Genbank Accession AXK59213.1) are included for reference. Amino acids are shaded to indicate shared identity among peptides.

### Enterovirus VP1 ELISA confirms VirScan Findings

Consistent with the VirScan data, the mean EV VP1 ELISA signal in AFM (n = 26, mean OD 0.51 IQR 0.56) was significantly higher than OND controls (n = 50, mean OD 0.08 IQR 0.06, p-value < 0.001 by Mann-Whitney test, Figure 3 and Supplemental Table 7). Mean EV signal detected by phage and ELISA demonstrated a linear correlation (R^2^ = 0.48, p-value < 0.001, Supplemental Figure 2). Among AFM patients, mean CSF EV antibody detected by either ELISA or VirScan did not differ based on whether EV RNA had been previously detected (n=15) or not (n = 11, mean OD 0.41 versus 0.65 by ELISA; mean rpK 6,444 vs 13,975 by VirScan, p-value = not significant for both comparisons). We attempted to identify whether a patient was infected with either EV-A71 or EV-D68 using both VirScan and ELISA but both assays yielded cross-reactivity, an expected issue with EV ELISA (Supplemental Figure 3). We did not observe an obvious independent effect of geography, year, or season on either the VirScan total EV or the ELISA VP1 EV data (Supplemental Figures 4-6).

**Figure 3.**
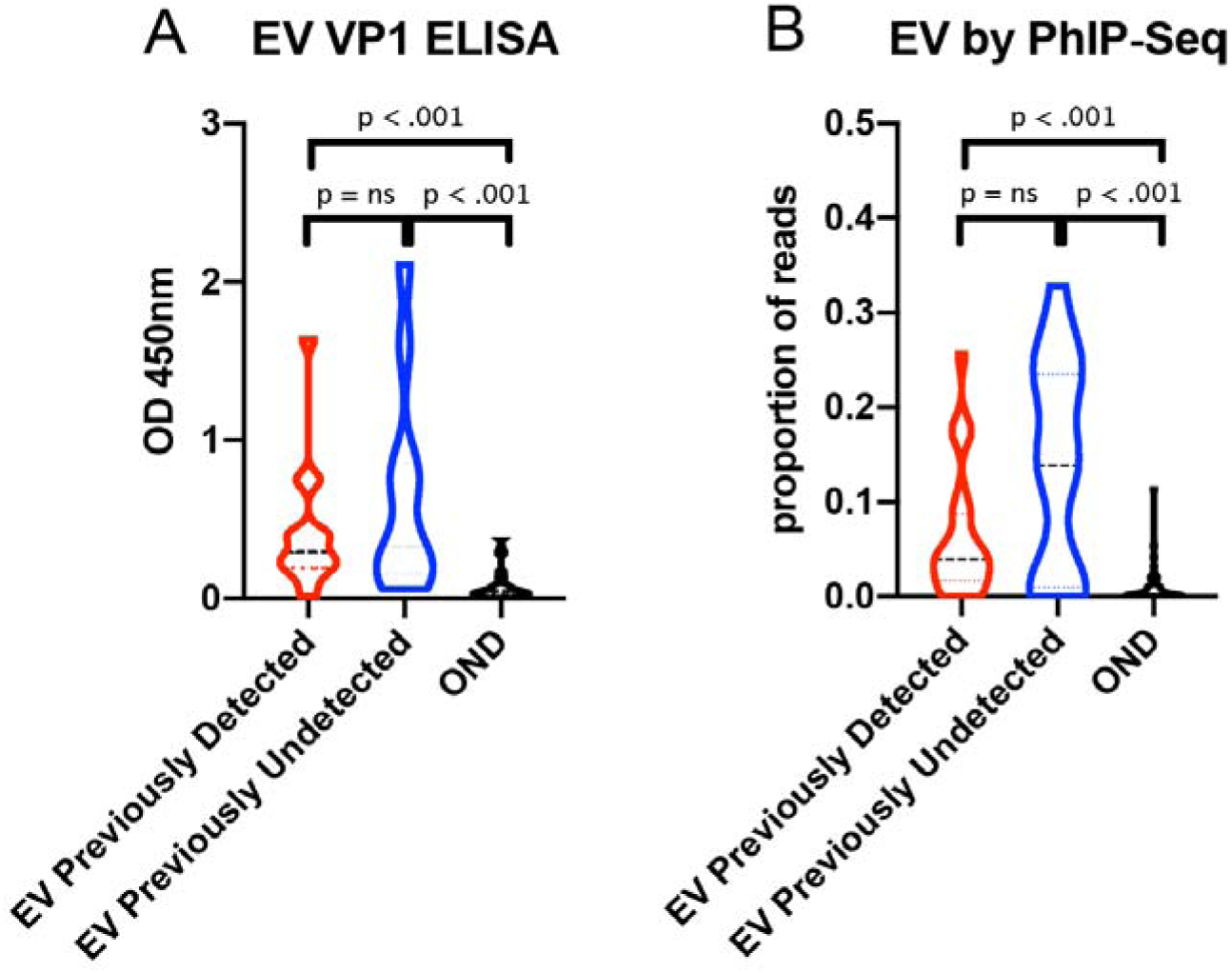
Independent Validation of Pan-Viral Phage Display with Purified Enterovirus VP1 Capsid Protein. **(A)** Violin plot that enterovirus (EV) signal generated by ELISA can be found at similar levels in patients with both previously detected (n=15) and previously undetected (n=11) EV infections (p = ns). In both AFM cohorts, there was a significantly greater amount of signal generated by ELISA compared with pediatric other neurologic disease (OND) controls (n=50) (p < 0.001 for both comparisons, Mann-Whitney test). (B) Similar results by VirScan with no differences seen when comparing EV signal in those with previously detected (n=23) and previously undetected (n=19) EV infection (p = ns). When each group was compared to the OND controls (n=58), both demonstrated significant enrichment of EV signal (p < 0.001; Mann-Whitney signed-rank test with Bonferroni adjustment for multiple comparisons).

## Discussion

We combined unbiased ultra-deep mNGS with an adaption of the VirScan method^17^ for comprehensively detecting anti-viral antibodies to query CSF from a relatively large (n=42) and geographically diverse subset of children presenting with AFM since 2014^17^. Ultra-deep mNGS combined with FLASH enrichment for EV-A71 and EV-D68 confirmed the presence of EV RNA in a single sample that was previously known to be positive for EV-A71 by PCR and failed to discover any other pathogen. There are a number of possible reasons for the lack of detectable EV nucleic acid in the CSF of AFM patients by mNGS or other methods. Clinically, radiologically and similarly to poliomyelitis, the CNS tissue involved in AFM is often restricted to the anterior horn cells in the cervical spinal cord, making it possible that little to no virus is shed into the CSF. In addition, children with AFM typically present with neurologic symptoms a median of 5-7 days after prodromal illness onset, decreasing the probability of RNA detection.^29^

Lack of consistent identification of viral nucleic acid in CSF is not limited to AFM, rather it is common to a wide range of neuroinvasive viruses, including poliovirus, rabies, West Nile virus, and other arboviruses.^30^ As a result, detection of intrathecal antibody production through CSF serologic testing is the gold standard for diagnosis of many neuroinvasive viruses, notably West Nile virus and varicella zoster virus.^31,32^ Thus, we supplemented CSF mNGS with VirScan to comprehensively profile CSF anti-viral antibodies in AFM cases and OND controls. VirScan revealed high levels of CSF immunoreactivity to immunodominant EV epitopes in AFM, independent of whether EV RNA had previously been detected in clinical testing of CSF or nonsterile sites. Independent testing with EV-A71 and EV-D68 VP1 ELISAs confirmed these findings. VirScan and whole VP1 ELISA were not able to consistently identify specific binding to individual EV types, likely owing to cross-reactive immune responses to conserved, linear EV antigens.^33^ There was a non-significant trend towards greater enrichment of EV antibodies in patients without directly detectable virus in a peripheral site, possibly owing to the rise in titer that occurs in the weeks following an infection.

This study has important limitations. First, detection of a serologic response to a virus at a single time point by itself does not fulfill Koch’s postulates for establishing causality between a virus and a particular disease. However, these serologic data support the specificity of the CSF antibody response to EVs in AFM, helping fulfill the Bradford Hill criteria for making a causal association.^13,14^ Second, further work will be necessary to establish, in a prospective manner, the diagnostic sensitivity and specificity for CSF EV serology, and thus we have only reported population level means and medians for our results. Third, our cases and controls were not optimally matched. Controls had OND, but case and control populations were not similar by age, year, or season, which are important risk factors for enterovirus infection in the United States. However, we did not see a significant effect of year or season on EV signal by VirScan or ELISA in the OND controls. Fourth, we did detect a statistically increased amount of signal to *Caliciviridae* in the AFM cases. However, the magnitude of signal was much less than for *Picornaviridae* and *Enterovirus*, and its clinical relevance is unclear as this family of viruses is not typically associated with neuroinvasive disease. We chose to report these preliminary findings, despite the limitations of the study design, because of the public health urgency of understanding the etiology of AFM. A prospective study with matched cases and controls is necessary to confirm our findings.

AFM is a potentially devastating neurologic syndrome whose incidence of reported cases has risen in the US since 2014 with biennial peaks. In addition, cases have now been detected in 14 other countries across 6 continents.^29^ There are no proven treatments for AFM, and like poliomyelitis, a vaccine may ultimately be the most effective prevention strategy. However, it is important to first achieve consensus around the likely etiologic agents. While continued vigilance for other possible etiologies of AFM is warranted, together, our combined mNGS, pan-viral VirScan, and viral protein ELISA interrogation of AFM CSF supports the notion that EV infection likely underlies the majority of AFM cases tested in this study. These results offer a roadmap for rapid development of EV CSF antibody assays to enable efficient clinical diagnosis of EV-associated AFM in the future.

## Acknowledgments

This work is supported by a National Multiple Sclerosis Society-American Brain Foundation Clinician Scientist Development Award FAN-1608-25607 (RDS), NIH grants K08NS096117 (MRW) and K23AI28069 (KM), the Chan Zuckerberg Biohub (JLD, CMT, JEP, EDC, WW, CKC, AL, MT, RS), an endowment from the Rachleff family (MRW), and the Sandler and William K. Bowes, Jr. Foundations (MRW, KCZ, HAS, CYC, LMK, and JLD). We would like to thank the patients and their families for their participation in this study.

## Disclaimer

The findings and conclusions in this report are those of the author(s) and do not necessarily represent the official position of the Centers for Disease Control and Prevention, the National Institutes of Health or the California Department of Public Health.

