## Supplemental Appendix for "Serological and metagenomic interrogation of cerebrospinal fluid implicates enteroviruses in pediatric acute flaccid myelitis"

**SUPPLEMENTARY APPENDIX**

**TABLE OF CONTENTS**

**List of Investigators**....................................................................................................................2-3

**Supplementary Methods**

**Patient Enrollment and Data collection**...........................................................................4

**Metagenomic Sequencing / FLASH**

Library Preparation...............................................................................................4-5

Pathogen Identification with IDseq......................................................................5-6

Immunoprecipitation of phage-bound patient antibodies.....................................8-9

Next-generation sequencing library preparation of phage DNA......................10-12

Bioinformatic Analysis of PhIP-Seq Data........................................................12-13

**Independent Enterovirus Antibody Validation**

Molecular Cloning of EV-D68 and EV-A71 VP1 proteins...................................13

Mammalian Expression of EV-A71 and EV-D68 VP1 proteins......................13-14

Purification of EV-A71 and EV-D68 VP1 proteins.........................................14-15

Indirect ELISA of EV-A71 and EV-D68 VP1 proteins.........................................15

**Table S1.**.......................................................................................................................................16

**Figure S1.**......................................................................................................................................17

**Figure S2.**......................................................................................................................................18

**Figure S3.**......................................................................................................................................19

**Figure S4.**......................................................................................................................................20

**Figure S5.**......................................................................................................................................21

**Figure S6.**......................................................................................................................................22

**Tables S2-S7.**......................................................................................................................Attached

**References**.....................................................................................................................................23

**Serological and metagenomic interrogation of cerebrospinal fluid implicates enteroviruses in pediatric acute flaccid myelitis**

Ryan D. Schubert, MD^1,2^, Isobel Hawes, BS^1*^, Prashanth S. Ramachandran, MBBS^1,2*^, Akshaya Ramesh, PhD^1,2*^, Emily D. Crawford, PhD^3,4^, John E. Pak, PhD^3^, Wesley Wu, PhD^3^, Carly K. Cheung, BS^3^, Brian D. O’Donovan, PhD^5^, Cristina M. Tato, PhD^3^, Amy Lyden, BS^3^, Michelle Tan, BS^3^, Rene Sit, BA^3^, Gavin Sowa, BS^6^, Hannah A. Sample, BS^5^, Kelsey C. Zorn, MHS^5^, Debarko Banerji, BS^2^, Lillian M. Khan, BS^5^, Riley Bove, MD^1,2^, Stephen L. Hauser, MD^1,2^, Amy A. Gelfand, MD, MAS^1^, Bethany Johnson-Kerner, MD, PhD^1,2^, Kendall Nash, MD^1^, Kalpathy S. Krishnamoorthy, MD^7^, Tanuja Chitnis, MD^7,8^, Joy Z. Ding, MD^9^, Hugh J. McMillan, MD, MSc^9^, Charles Y. Chiu, MD, PhD^10^, Benjamin Briggs, MD, PhD^11^, Carol A. Glaser, DVM, MPVM, MD^12^, Cynthia Yen, MPH^13^, Victoria Chu, MD, MPH^13^, Debra A. Wadford, PhD^13^, Samuel R. Dominguez, MD, PhD^14^, Terry Fei Fan Ng, PhD^15^, Rachel L. Marine, PhD^15^, Adriana S. Lopez, MHS^15^, W. Allan Nix, BS^15^, Ariane Soldatos, MD, MPH^16^, Mark P. Gorman, MD^17^, Leslie Benson, MD^17^, Kevin Messacar, MD^14^, Jennifer L. Konopka-Anstadt, PhD^15^, M. Steven Oberste, PhD^15^, Joseph L. DeRisi, PhD^3,5^, Michael R. Wilson, MD, MAS^1,2**^

*Authors contributed equally

Affiliations:

^1^UCSF Weill Institute for Neurosciences, San Francisco, CA, USA

^2^UCSF Department of Neurology, San Francisco, CA, USA

^3^Chan Zuckerberg Biohub, San Francisco, CA, USA

^4^UCSF Department of Microbiology and Immunology, San Francisco, CA, USA

^5^UCSF Department of Biochemistry & Biophysics, San Francisco, CA, USA

^6^UCSF School of Medicine, San Francisco, CA, USA

^7^Department of Neurology, Massachusetts General Hospital, Boston, MA, USA

^8^Department of Neurology, Brigham and Women's Hospital, Boston, MA, USA

^9^Division of Neurology, Children's Hospital of Eastern Ontario, University of Ottawa, Ottawa, Ontario, Canada.

^10^Department of Laboratory Medicine and Medicine, Division of Infectious Diseases, University of California, San Francisco, San Francisco, USA

^11^Department of Pediatrics, Division of Infectious Diseases, University of California, San Francisco, San Francisco, USA

^12^Department of Pediatric Infectious Diseases, Kaiser Permanente Oakland Medical Center, Oakland, California, USA

^13^Division of Communicable Disease Control, California Department of Public Health, Richmond, California

^14^Children’s Hospital Colorado and Department of Pediatrics, University of Colorado School of Medicine, Aurora, CO, USA.

^15^Division of Viral Diseases, Centers for Disease Control and Prevention, Atlanta, GA, USA

^16^National Institute of Neurological Disorders and Stroke (NINDS), NIH, Bethesda, Maryland, USA.

^17^Department of Neurology, Boston Children’s Hospital, Boston, MA, USA

**SUPPLEMENTAL METHODS**

**Patient Enrollment and Data Collection**

Inclusion criteria for patients with a confirmed diagnosis of definite or probable acute flaccid myelitis are outlined by the Council of State and Territorial Epidemiologists (Supplemental Table 1).^1^ All patients were evaluated by board-certified neurologists at their respective enrollment sites or at the Centers for Disease Control and Prevention (CDC). Patients 1 through 8 were enrolled at Boston Children’s Hospital. Patients 25 through 28, and 47 through 100 were enrolled in a research study at UCSF (IRB number 13-12236) for pathogen and autoantibody detection for patients with idiopathic neuroinflammation. Patients 9 through 17 were enrolled through the University of Colorado (IRB number 12-0745). Patients 18 through 24 were from the California Department of Public Health (CDPH). Patients 29-46 were from the Division of Viral Diseases at the CDC, with this work constituting public health surveillance as determined by the human subjects coordinator of the National Center for Immunization and Respiratory Disease. Electronic and paper medical records of patients from Boston Children’s Hospital, UCSF, the University of Colorado, and the CDPH were reviewed for demographic details, clinical data, laboratory results, and outcome at last follow-up. Age and sex were shared in aggregate from the CDC.

**Metagenomic Sequencing/FLASH**

**Library Preparation**

RNA was extracted from 100 µl of CSF using the Zymo ZR Duet DNA/RNA MiniPrep Plus kit (Zymo Cat. No. D7003). RNA was also extracted from a non-templated control (water) for quality control. Sequencing libraries were prepared with New England Biolabs’ NEBNext Ultra II RNA Library Prep Kit for Illumina (Cat. No. E7770) on the Labcyte Echo 525 and the Biomek NX MC using a previously described protocol, optimized and adapted for miniaturization and automation.^2^ External RNA Controls Consortium (ERCC) spike-ins were used to back calculate RNA mass input. The mNGS libraries were pooled using the Labcyte Echo 525 based on preliminary sequencing on the Illumina iSeq. Host ribosomal RNA was then depleted from the pooled library using Depletion of Abundant Sequences by Hybridization (DASH).^3^ Final sequencing was performed on the NovaSeq 6000 to generate 150 nucleotide (nt), paired-end reads.

**FLASH-NGS Sequencing Library Preparation**

A library of Cas9 guide RNAs suitable for FLASH-NGS was designed using the FLASHit software. Input sequences consisted of 39 EV-D68 and 8 EV-A71 sequence clusters derived from 100% Cluster Database at High Identity with Tolerance (CD-HIT) of all US EVD68 sequences from the 2014 outbreak and 9 sequences from the 2016 Catalonia EVA71 (Supplemental Table 2). Guide RNAs from these sequences were transcribed from DNA templates (Integrated DNA Technologies) as described.^4^ RNA from AFM and OND cases was reverse transcribed using the First and Second Strand Synthesis modules from the NEBNExt Ultra II RNA Library Prep Kit. Resulting cDNA was then subjected to FLASH as previously described. Libraries were sequenced on a NextSeq 550 instrument to generate 150 nt, paired-end reads. A mean of 2.7 million reads per sample were collected (range 0.28 million to 8.6 million).

**Pathogen Identification**

Microbial pathogens were identified from raw sequencing reads using the IDseq (v3.2) Portal (https://idseq.net), a cloud-based, open-source bioinformatics platform designed for detection of microbes from metagenomic data. To distinguish potential pathogens from ubiquitous environmental agents including laboratory reagent contaminants and skin commensal flora, a Z-score was calculated for both nucleic acid and protein alignments for each genus, relative to a background of controls. FLASH reads were aligned to the reference sequences in Supplemental Table 2 using bowtie2 with the --very-sensitive flag.^5^

**FLASH Enrichment**

FLASH enrichment was calculated by dividing the number of reads detected as a proportion of all reads sequenced in the FLASH library by that found in the standard mNGS libraries. For the purpose of FLASH enrichment, we re-aligned the mNGS reads to the FLASH reference sequences to prevent over-inflation that could result from CD-HIT-DUP in the IDseq pipeline.^4^

**Calculation of RNA mass input with ERCC**

ERCCs were spiked into all samples at a predetermined mass of 25pg (ThermoFisher Scientific). STAR alignment within the IDseq pipeline provided total ERCC reads and read counts for each individual ERCC sequence. Total RNA mass input is calculated using the following ratio: (ERCC mass input/Total mass input) = (ERCC sequencing reads/Total sequencing reads). ERCC mass was removed from total RNA mass to provide the original RNA input mass. The inputted mass for each individual ERCC strand was calculated from the manufacturers prespecified concentrations. Individual ERCC sequences were then ranked based on mass. Detection of the ERCC sequence with the lowest mass was used to determine the sequencing limit of detection.

**Pan-Viral Phage Display with Immunoprecipitation (PhIP-Seq)**

**Library Design**

We developed a custom viral phage display library of 481,966 62 amino acid peptides with a 14 amino acid overlap. These overlapping oligonucleotide sequences were tiled across the protein sequences of all viruses downloaded from the UniProtKB and National Center for Biotechnology Information’s RefSeq databases in February 2017. Viruses exclusively from these hosts were removed: algae, archaea, bacteria, diatom, environment, fungi, plant. Additionally, all giant virus genomes (>1 Gb) were removed. The final set of viruses were clustered using CD-HIT at 98% at the amino acid level except for overrepresented viruses which were de-clustered on the following percent identity: HIV/SIV (92%), hepatitis B virus (95%), porcine reproductive and respiratory viruses (95%) and human papillomavirus (97%). Additionally, clinically relevant viruses, not included in these databases, were manually added (Jamestown Canyon virus, La Crosse virus, Cache Valley virus, Argentinian hemorrhagic fever virus, Bolivian hemorrhagic fever virus, Venezuelan hemorrhagic fever virus, human herpes virus 5, California encephalitis virus, bourbon virus, smallpox virus, herpesvirus B, enterovirus D68, coltivirus, rubulavirus, human respirovirus, pteromalus puparum peropuvirus, human cosavirus F1, cardiovirus, and Vilyuisk human encephalomyelitis virus). The following vaccine strains were added: measles, mumps, polio, rubella, and influenza viruses (vaccine strains between 2012-2017). Technical controls were included from six other genes (glutamate ionotropic receptor NMDA type subunit 1 (GRIN1), glial fibrillary acidic protein (GFAP), Gephyrin, green fluorescent protein, MYC Proto-Oncogene, and tubulin) were also added.

Viral peptide sequences included were again clustered at 95% nucleotide identity using CD-HIT. Next, low-complexity sequences were removed with the Lempel-Ziv-Welch algorithm (LZW score < 0.04). Additionally, sequences with homopolymer runs greater than 20 amino acids were removed. To minimize self-priming of oligonucleotides during PCR, synonymous sequence scrambling by swapping with a synonymous codon picked at random in the overlapping region of the peptide, and restriction site (EcoR1, HindIII and XhoI) removal, was performed. A 5’ mixed alanine glycine linker and 3’ region with two stop codons were added (5'=GTAGCTGGAGTTGTTGCAGGC 3'=TGATAAGCATATGCCATGGCCTC). The final set of oligonucleotides was synthesized by Agilent. Libraries were cloned, packaged, amplified, and stored in an 8% glycerol containing media according to the manufacturer’s instructions (T7 Select 10-3 Cloning Kit, EMD Millipore).

**Preparation of phage libraries from stocks**

Phage libraries were prepared and amplified fresh from 80°Cfrozen glycerol stocks. To prepare phage libraries, a 500 mL culture of *E. coli* BLT5403 was incubated at 37^o^C with shaking until mid-log phase, defined as OD_600_= 0.5. The culture was then inoculated at a multiplicity of infection (MOI) of 0.001 and lysed at 37°C for 3 hours. Lysate was spun at 3500 g for 15 minutes to remove debris. Phage-containing supernatant was then precipitated with PEG/NaCl (PEG-8000 20%, NaCl 2.5 mM) overnight at 4°C. Precipitated phage were then pelleted for 15 minutes at 10,000 g at 4°C and re-suspended in storage media (20 mM Tris-HCl, pH 7.5, 100 mM NaCl, 6 mM MgCl_2_) before 0.22uM filtration. Resulting phage libraries were titered by plaque assay and adjusted to a working concentration of 10^10^-10^11^ pfu/mL.

**Immunoprecipitation of phage-bound patient antibodies**

Liquid handling steps were carried out on the Biomek FX robotics platform (Beckman Coulter) as previously described.^6^ 96-well hardshell PCR plates (Biorad) were blocked overnight with blocking buffer (3% immunoglobulin-depleted BSA, PBS-T) to prevent nonspecific binding of phage to walls of the plate. Blocking buffer was replaced with 150 µL of phage library. Patient samples were diluted 1:1 with storage buffer (final concentration 20% glycerol, 20mM HEPES pH 7.3, 0.02% NaN_3_ in DPBS) and stored in pre-blocked 96-well plates for use during an experiment. Two microliters of CSF diluted in storage buffer were incubated with phage library overnight on an orbital shaker at 4°C.

Pierce Protein A/G magnetic beads (ThermoFisher Scientific) were washed by magnetic separation and re-suspended in an equivalent volume of 0.1% TNP-40 (150 mM NaCl, 50 mM Tris-HCl, 0.1% NP-40) prior to use. Ten microliters of A/G bead slurry was added to each phage-CSF mixture and immunoprecipitated at 4°C on an orbital shaker for 1 hour to allow patient antibodies to bind to beads. The beads were then washed three times by magnetic separation with 150 µL 0.1% TNP-40. Subsequent to the final wash, beads were re-suspended in 150 µL of Luria broth (LB) and used to inoculate 96-deep-well plates containing 1 mL of fresh mid-log *E. coli* cultures (OD_600_= 0.5). Cultures were lysed for 3 hours at 37°C with a shaker (Infors) and then centrifuged for 10 minutes at 3500 g to remove debris. One hundred fifty microliters of lysate were then mixed with an additional 2 µL of patient sample overnight 4°C and immunoprecipitation was repeated to further enrich for relevant antibody-binding phage. Lysates were freshly frozen at -20°C for storage before sequencing library preparation.

**Next-generation sequencing library preparation of phage DNA**

Plates containing lysates from both IP1 and IP2 were thawed and subsequently diluted 1:50 in ddH_2_O. Enrichment of phage DNA and barcoding of individual IP reactions was performed using a single PCR reaction using nested PCR with the following reagents (ThermoFisher Scientific):

| Reagents | µL/reaction |
| --- | --- |
| 5x HF buffer | 5 |
| DMSO | 0.75 |
| 10 mM dNTPs | 0.5 |
| Phusion polymerase | 0.25 |
| Water | 16.25 |
| Pan-OME primer mix (see below) | 1.25 |
| 5 µM TruSeq multiplexing primers | 2.5 |
| DNA template (1:50) | 1 |

*pan-OME primer mix:*

panOME_rev0:

GTGACTGGAGTTCAGACGTGTGCTCTTCCGATCTTAGTTACTCGAGTGCGGCCGCAAGC

panOME_rev1:

GTGACTGGAGTTCAGACGTGTGCTCTTCCGATCTNTAGTTACTCGAGTGCGGCCGCAAGC

panOME_rev2:

GTGACTGGAGTTCAGACGTGTGCTCTTCCGATCTNNTAGTTACTCGAGTGCGGCCGCAAGC

panOME_rev3:

GTGACTGGAGTTCAGACGTGTGCTCTTCCGATCTNNNTAGTTACTCGAGTGCGGCCGCAAGC

panOME_forward

ACACTCTTTCCCTACACGACGCTCTTCCGATCTAGTCAGGTGTGATGCTCGGGGATCC

*Primer design for Truseq multiplexing primers*

Forward:

AATGATACGGCGACCACCGAGATCTACAC[NNNNNNN]GGAGCTGTCGTATTCCAGTCAGGTGTGATGCTC

NNNNNNN = i5 index

Reverse:

CAAGCAGAAGACGGCATACGAGAT[NNNNNNN]GGTTAACTAGTTACTCGAGTGCGGCCGCAAGC

NNNNNNN = i7 index

*Thermocycling conditions:*

95°C for 2 minutes

5 cycles of: 95°C x30 sec, 63°C x30 sec, 72°C x45 sec

5 cycles of: 95°C x30sec, 60°C x30sec, 72°C x45sec

15 cycles of: 95°C x30sec, 58°C x30sec, 72°C x45sec

72°C for 2 minutes

hold at 10°C

After the PCR reaction completed, products were pooled (3 µL per sample) and cleaned with a 1x bead clean-up (SPRISelect, Beckman Coulter). Product concentration was quantified using an automated electrophoresis system (Bioanalyzer High-Sensitivity DNA, Agilent). Libraries were sequenced on the HiSeq 4000 (Illumina) using 150 nt paired-end reads.

**Bioinformatic Analysis of PhIP-Seq Data**

Reads were quality-filtered, paired-end reconciled (PEAR v0.9.8), and aligned to a reference database of the full library (bowtie2 v2.3.1).^5^ SAM files were parsed using custom analysis tools (Python/Pandas) and individual phage counts were normalized to reads per 100,000 (rpK) by dividing by the sum of counts and multiplying by 100,000 as previously described.^6^ Peptide-level fold changes were calculated by dividing individual peptide rpK values with the mean rpK value detected for the peptide in null immunoprecipitations (i.e. A/G beads alone, n = 4 replicates) and taking the log_10_ of the result. For peptides not observed in the control IP, a value of 1 rpK was assumed to calculate the log_10_ fold change. During downstream PhIP-Seq analysis, we only considered phage with fold change values 10-fold above the null background. To account for multiple tests, we adjusted p-values with the Bonferroni procedure.

**Independent Enterovirus Antibody Validation**

**Molecular Cloning of EV-D68 and EV-A71 VP1 proteins:**

The nucleotide sequences for VP1 from EV-D68 (AIT52326) and EV-A71 (MH718269), modified to include an N-terminal start codon (Met) and a C-terminal (GGGGS)_3_ linker followed by a hexa-histidine tag (his-tag), were codon optimized, synthesized, and cloned in between the EcoRI-NheI sites of the mammalian expression plasmid pTwist CMV BetaGlobin (Twist Biosciences), followed by plasmid scale up (Molecular Cloning Laboratories).

**Mammalian Expression of EV-A71 and EV-D68 VP1 proteins:**

Suspension ExpiCHO cells, maintained at 37°C, 120 RPM (25 mm shaking diameter), 8% CO_2_, and 80% humidity, were transiently transfected with pTwist CMV BetaGlobin VP1 expression plasmids at 30 mL scale in 150 mL polycarbonate, disposable, sterile vented Erlenmeyer shake flasks (Corning), following the manufacturer’s guidelines. One day before transfection, cells were split to a final density of 4 million viable cells/mL. On the day of transfection, cells at 98% viability were split to a final density of 6 million viable cells/mL and transfected using ExpiFectamine and 30 μg of plasmid DNA. 20 hours after transfection, 300 μl of ExpiCHO enhancer and 12 mL of ExpiCHO feed was added to each flask, followed by shifting the cells to 32°C, 120 RPM (25 mm shaking diameter), 5% CO_2_, 80% humidity. Six days after transfection, we observed >50% cell viability (>10 million total cells/mL), measured using a Countess II FL automated cell counter (Thermo Fisher Scientific). Cells were harvested at this time by centrifugation at 4°C, 500 g for 5 minutes, washed with 1X DPBS and frozen at -80°C.

**Purification of EV-A71 and EV-D68 VP1 proteins:**

Harvested cells were thawed and resuspended in a lysis buffer consisting of 1% Triton X-100 in 1X PBS buffer, 1X EDTA-free cOmplete protease inhibitor cocktail (Sigma Aldrich) at a ratio of 1 mL lysis buffer per 10 million total cells.  Resuspended cells were mixed with end-over-end mixing at 4°C for 1 h, centrifuged at 14,000 g for 10 minutes, and the supernatant (i.e. clarified lysate) was collected. Clarified lysate was concentrated using Amicon Ultra centrifugal filters (Millipore-Sigma, 10 kDa MWCO) to a final volume below 10 mL. Imidazole (Teknova, 2 M solution, pH 7.5) was added to the concentrated lysate to a final concentration of 10 mM.  0.5 mL of His-Pur Ni-NTA Resin (ThermoFisher Scientific) was pre-equilibrated in 1 mL of equilibration buffer (PBS, 1% Triton X-100, 10 mM imidazole, pH 7.4) and then added to the concentrated lysate in a 50 mL conical tube. All subsequent capture, wash, and elution steps described below were carried out with end-over-end mixing at 4°C followed by centrifugation (700 g, 2 min, 4 °C) after each step to separate the resin from the supernatant.  Following incubation for 1 h to allow for VP1 capture, the Ni-NTA resin was washed with 6 x 25 mL wash buffer (PBS, 1% Triton X-100, 40 mM imidazole, pH 7.4) for 10 minutes per wash. After the final wash, the resin was transferred to a 1.5 mL microcentrifuge tube. Captured VP1 was eluted from the resin with 5 x 1 mL of elution buffer 1 (PBS, 1% Triton X-100, 250 mM imidazole, pH 7.4) followed by 1 x 1 mL of elution buffer 2 (PBS, 1% Triton X-100, 500 mM imidazole, pH 7.4). The elution fractions were pooled and desalted using PD10 columns (GE Healthcare) and concentrated to ~1 mL using Amicon Ultra centrifugal filters (10 kDa MWCO). VP1 protein titer was determined by anti-his Western blotting, using 4-15% mini-protean TGX stain-free protein gels (Bio-Rad) transferred to a PVDF membrane, a 6x-His Tag monoclonal primary antibody (HIS.H8, ThermoFisher Scientific) at 1:3000 dilution in PBS for 1 hr, an alkaline phosphatase secondary antibody (#31320, ThermoFisher Scientific) at 1:5000 dilution in PBS for 1hr, and Western Blue stabilized substrate for alkaline phosphatase (Promega) for detection. The standard curve for protein titer determination was generated using recombinant human his-tagged albumin (ProSci), loaded as a 5-point serial dilution from 50 to 3.125 ng.

**Indirect ELISA of EV-A71 and EV-D68 VP1 proteins:**

Nickel-coated 96-well polystyrene plates were coated overnight at 4°C with 50 µL 0.5-2 µg/mL of purified EV-D68 or EV-A71 VP1 diluted in PBS. Wells were washed 3 times with PBS-T using a Bio-Tek ELX 405 microplate washer. Wells were then blocked overnight at 4°C with 200 µL blocking buffer containing PBS, 2% immunoglobulin-depleted bovine serum albumin (Jackson ImmunoResearch Laboratories), and 0.05% Tween 20 (Sigma). Wells were again washed 3 times. CSF diluted 1:1 in storage buffer as described above was further diluted to a final concentration of 1:40 in blocking buffer, final volume 50 µL, and incubated overnight at 4°C. Wells were washed 6 times with 2 minute soaking intervals. Goat anti-human IgG (H+L) HRP (Invitrogen) was diluted 1:500 into 1/5 blocking buffer with PBS and 50 µL then added to each well for 1 hour at room temperature. Wells were washed again 6 times with 2 minute soaking intervals. Fifty microliters of 1-Step Ultra TMB-ELISA substrate solution was added to each well and the reaction stopped after 10 minutes with ELISA stop solution (ThermoFisher Scientific). Optical density (OD) was measured at 450 nm using a SpectraMax M5 Multi-Mode Microplate Reader (Molecular Devices) and background signal subtracted.

Supplemental Table 1 – Criteria for the diagnosis of AFM

**Clinical Criteria**

An illness with onset of acute flaccid myelitis

**Laboratory Criteria for Diagnosis**

Confirmatory Laboratory Evidence: a magnetic resonance image (MRI) showing spinal cord lesion largely restricted to gray matter*† and spanning one or more vertebral segments

Supportive Laboratory Evidence: cerebrospinal fluid (CSF) with pleocytosis (white blood cell count >5 cells/mm3)

**Case Classification**

**Probable**

Clinically compatible case AND

Supportive Laboratory Evidence: cerebrospinal fluid (CSF) with pleocytosis (white blood cell count >5 cells/mm3)

**Confirmed**

Clinically compatible case AND

Confirmatory laboratory evidence: MRI showing spinal cord lesion largely restricted to gray matter*† and spanning one or more spinal segments

* Spinal cord lesions may not be present on initial MRI; a negative or normal MRI performed within the first 72 hours after onset of limb weakness does not rule out AFM.

†Terms in the spinal cord MRI report such as “affecting mostly gray matter,” “affecting the anterior horn or anterior horn cells,” “affecting the central cord,” “anterior myelitis,” or “poliomyelitis” would all be consistent with this terminology.

Supplemental Figure 1 – Flow chart depicting patient enrollment by institution.

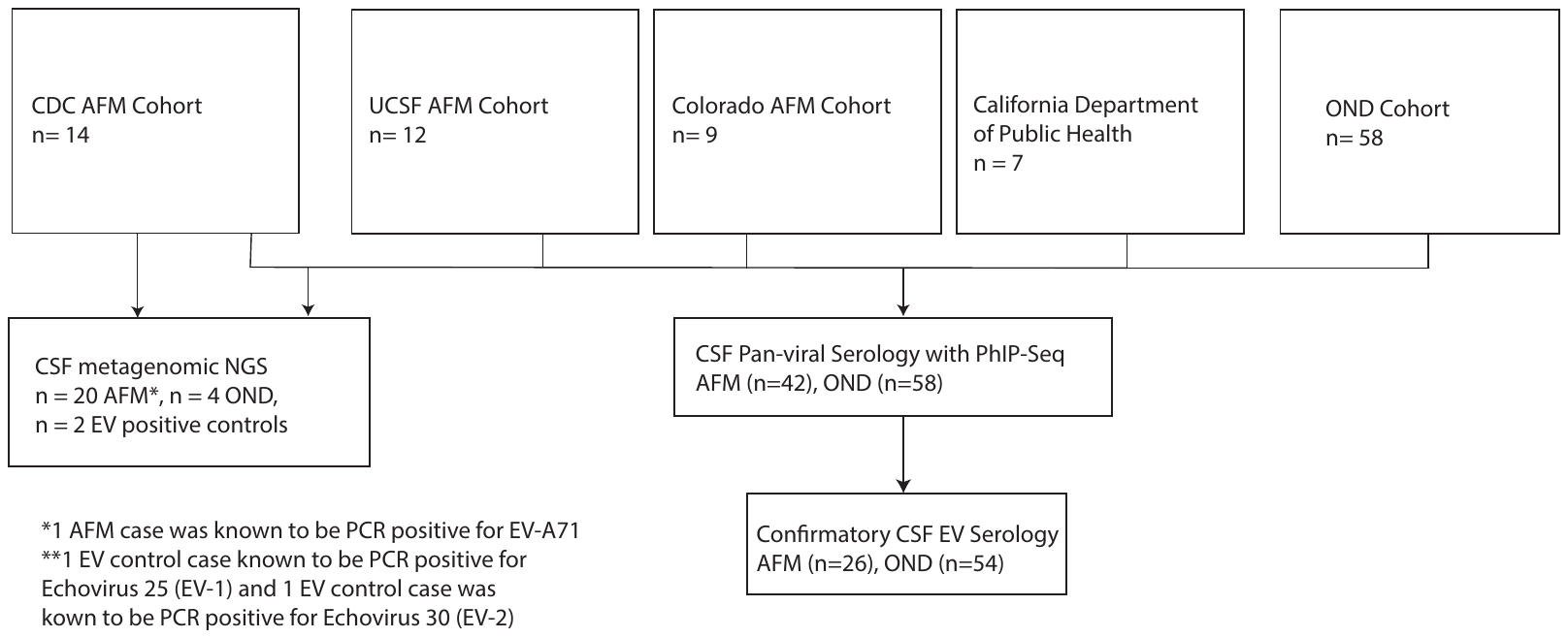

mNGS with and without FLASH were performed on CSF samples acquired from the CDC (n=14 AFM, n=4 OND, n=2 EV positive controls) and UCSF AFM Cohort (n=6 AFM). Samples from all institutions were tested by PhIP-Seq. Due to limited sample, a subset of those tested by PhIP-Seq were tested by confirmatory ELISA.

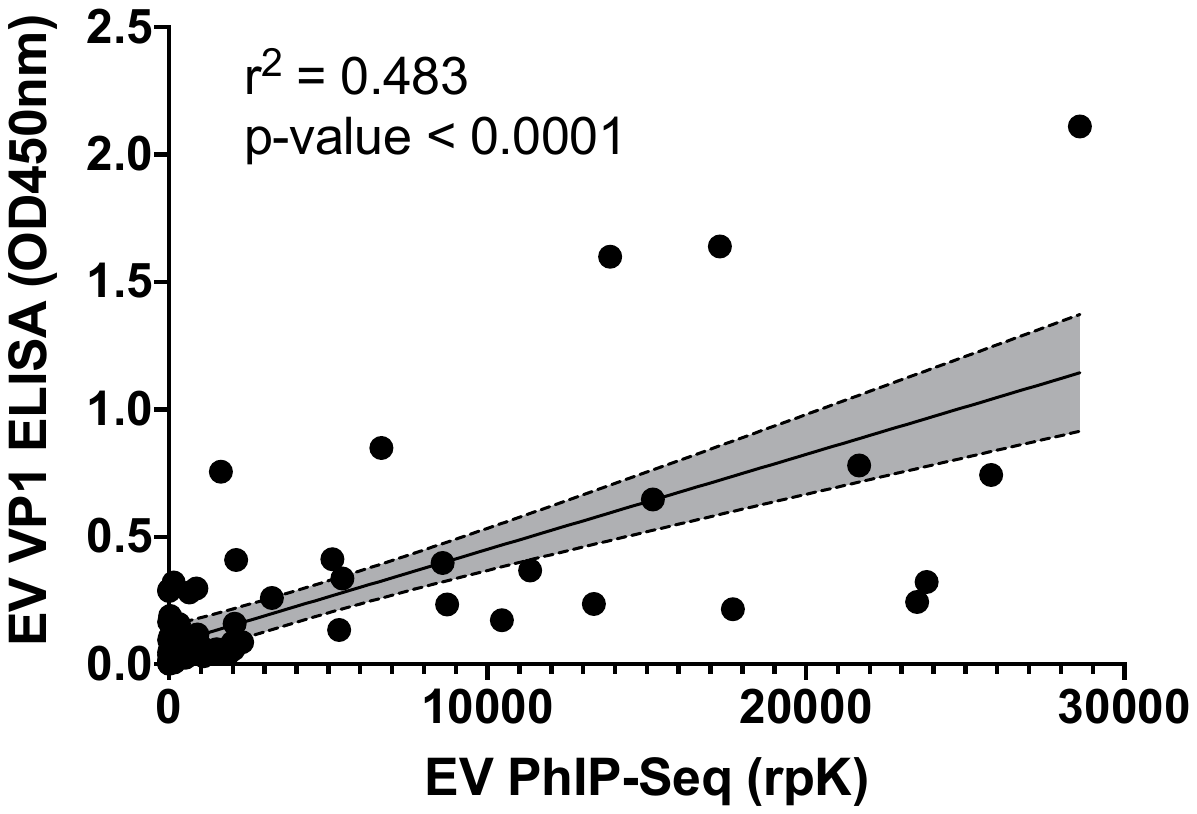

Supplemental Figure 2 – Comparison of PhIP-Seq and ELISA

A comparison of the total amount of enterovirus signal generated by PhIP-Seq (x-axis) to the maximum OD generated by either EV-D68 or EV-A71 signal ELISA (greater of the two values shown) for all samples run (n = 26 AFM + 50 OND). The 95% confidence intervals are shaded in grey.

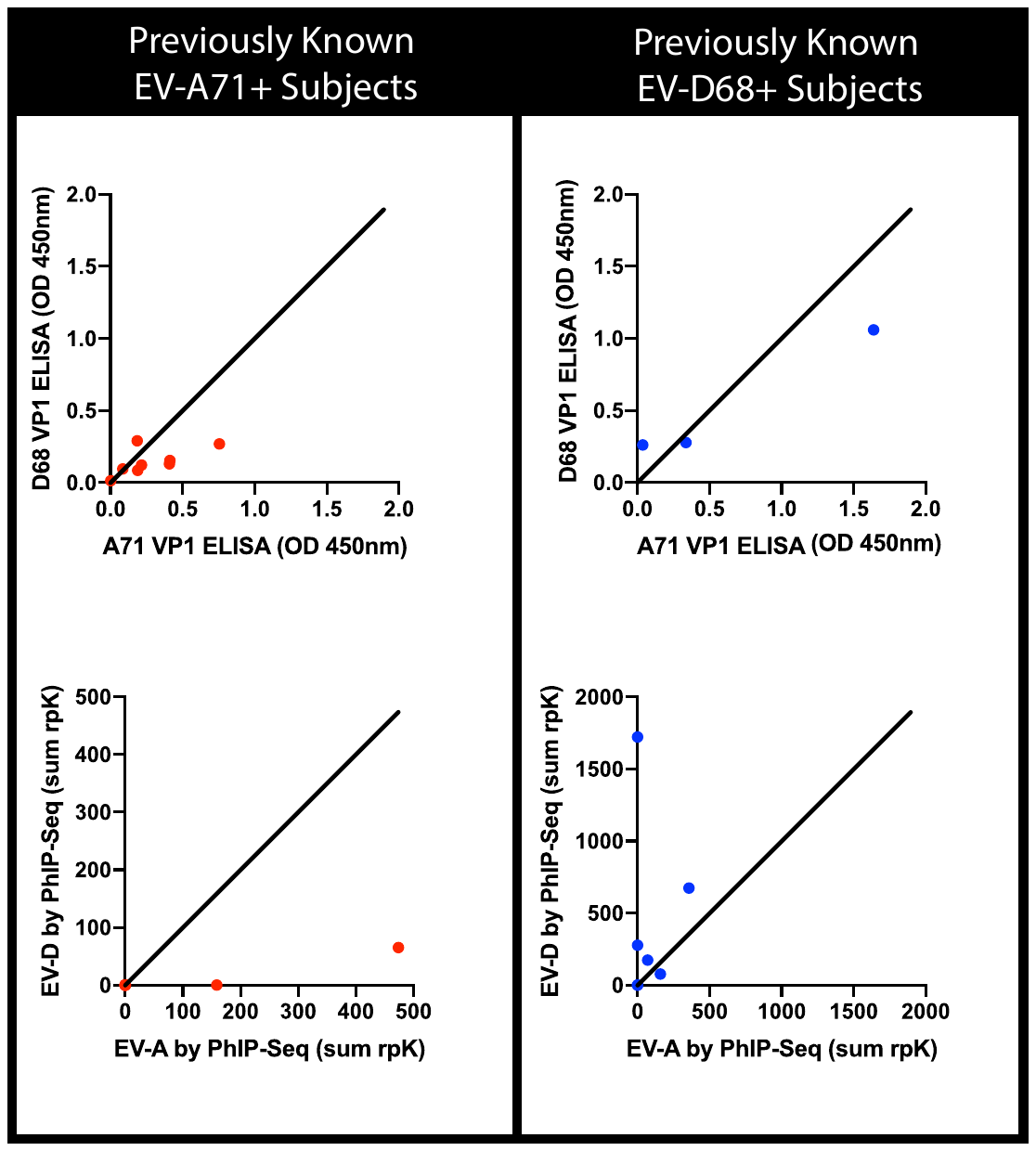

Supplemental Figure 3 – Strain calling by ELISA versus PhIP-Seq

ELISA and PhIP-Seq data from subjects with EV-A71 or EV-D68 detected by RT-PCR in either CSF, stool or respiratory fluid. Top panels with strain-specific VP1 ELISA data from EV-A71 (n = 8, red) and EV-D68 (n = 3, blue) patients show cross reactivity. Bottom panels shows PhIP-Seq data from known EV-A71 (n = 9, red) and EV-D68 (n = 7, blue) patients. EV-A and EV-D signals were generated by summing the total rpK generated against EV-A and EV-D derived peptides within a sample.

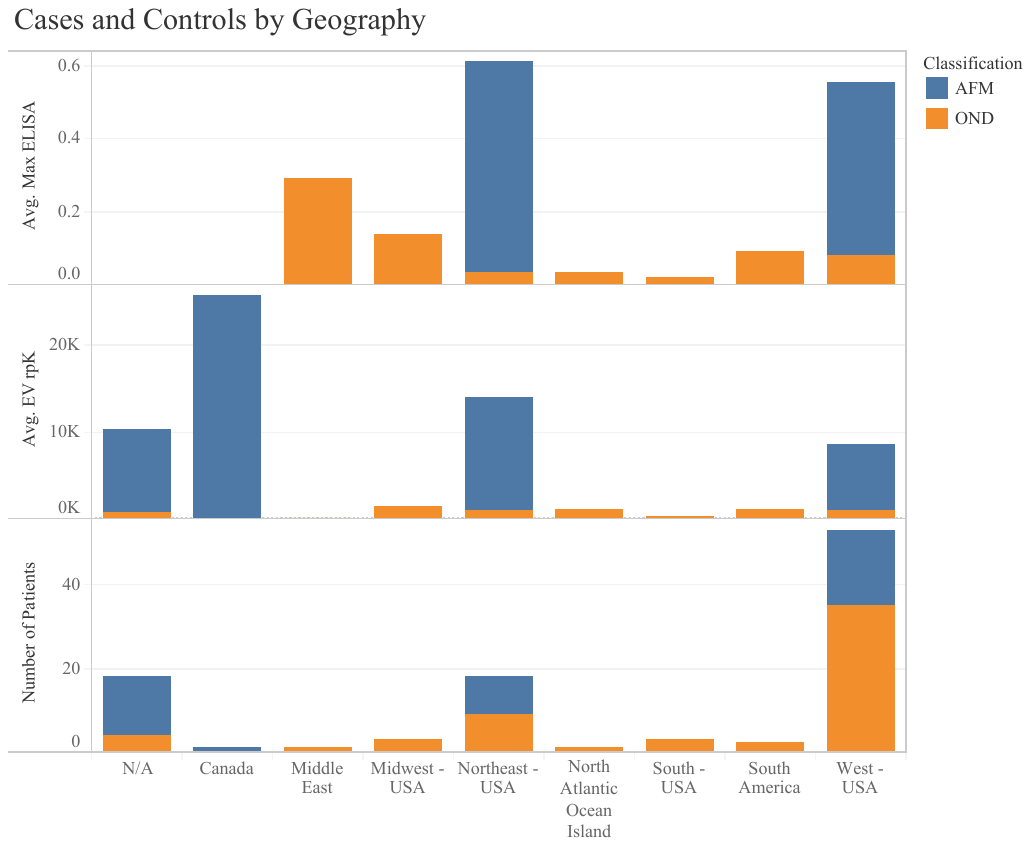

Supplemental Figure 4 – Geographic comparison of cases (blue) and controls (orange) with average EV signal by ELISA (top), average EV signal by PhIP-Seq (middle), and total number (bottom).

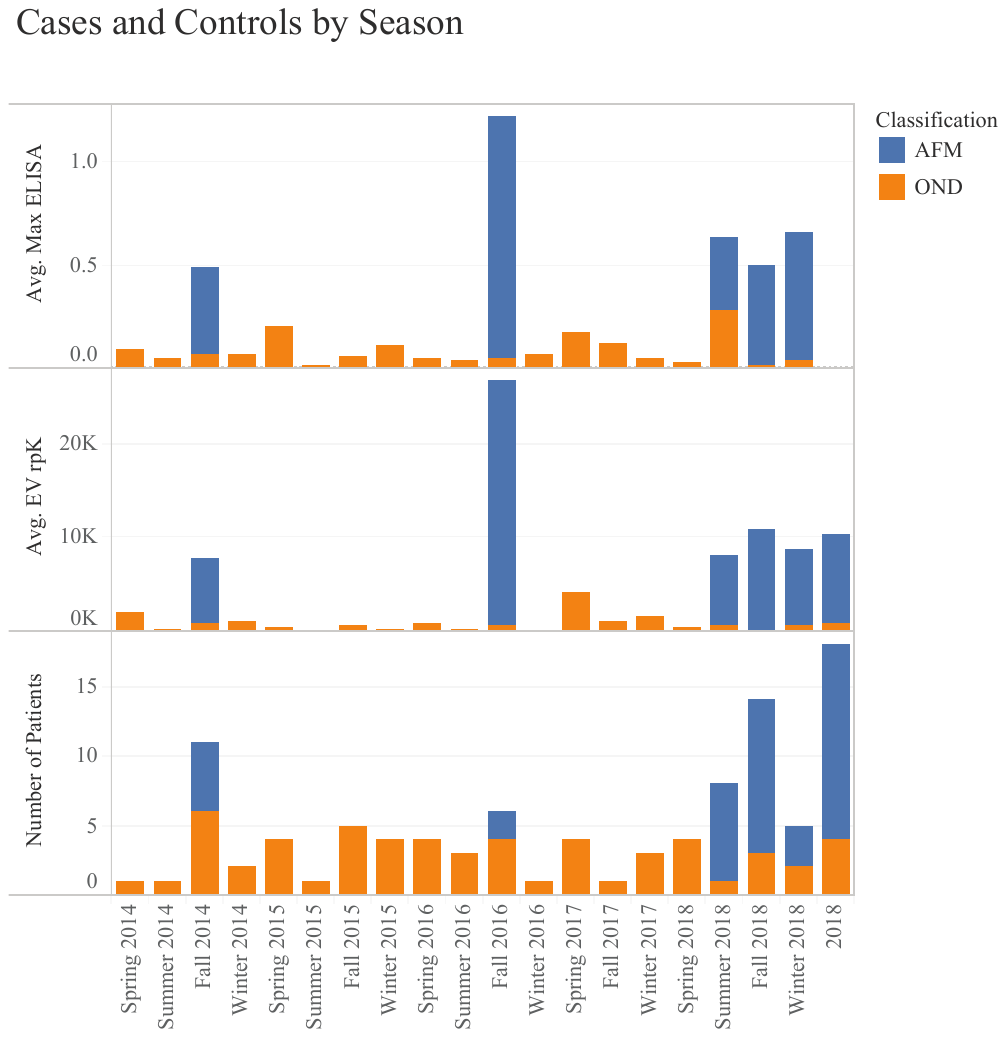
 Supplemental Figure 5 – Season and year comparison for cases (blue) and controls (orange) with average EV signal by ELISA (top), average EV signal by PhIP-Seq (middle), and total number (bottom).

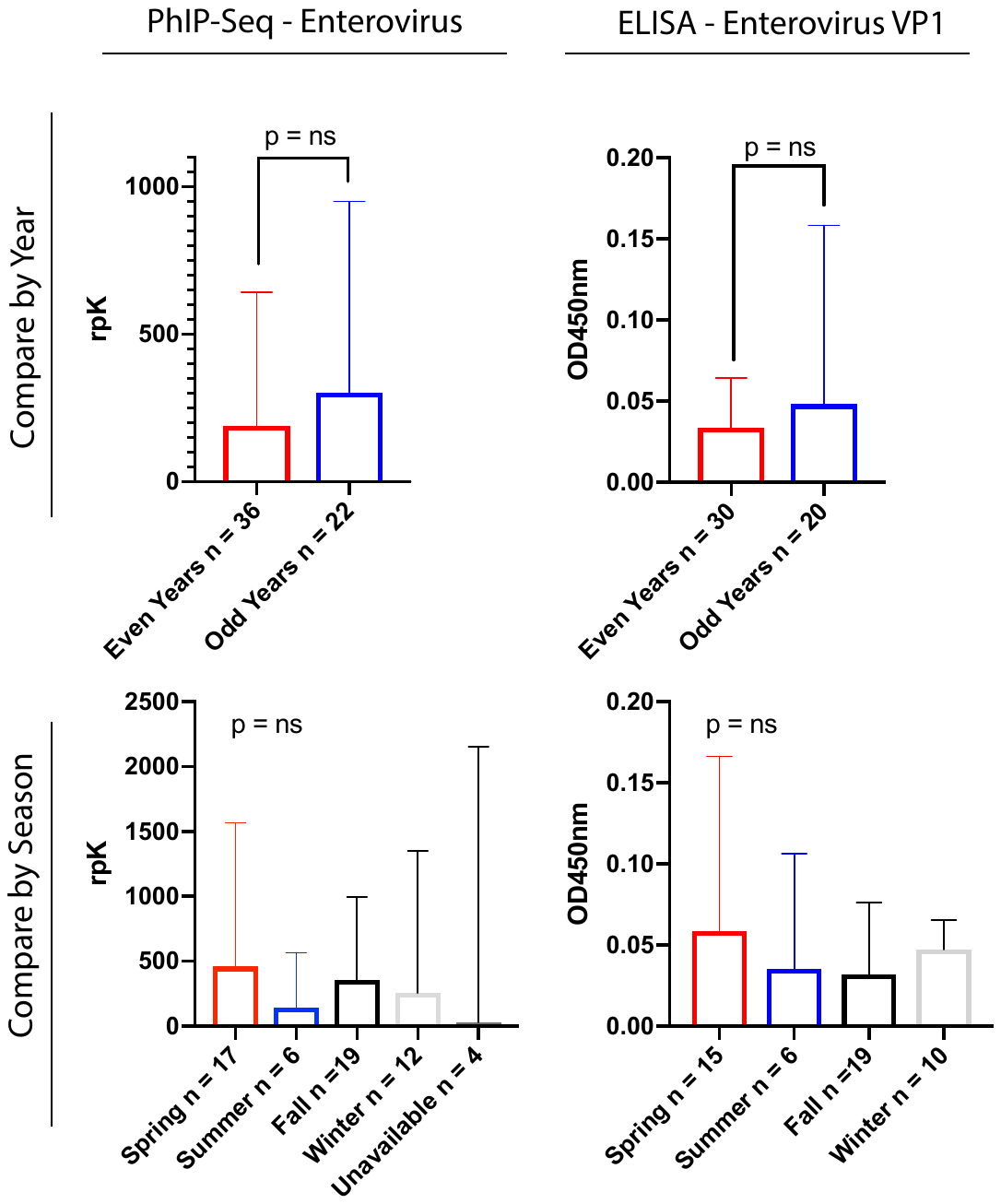

Supplemental Figure 6 – Analysis of effect of year and season on enterovirus signal in the OND controls. EV PhIP-Seq (left, n = 54) and EV VP1 ELISA (right, n = 50) for the OND control cohort by year (top) and season (bottom). Bar graphs depict heights as median values with error bars reflecting the interquartile range. Statistics for year were performed with the Mann-Whitney test and for seasons with the Kruskal-Wallace test.
